# Warming-induced range expansion effects on the diversity and composition of the gut microbiome: a case study with two butterflies

**DOI:** 10.1101/2023.01.11.523549

**Authors:** Daniel Montoya, Maaike de Jong

## Abstract

Species are the habitat of a diverse community of organisms, including bacteria, fungi and viruses – the *microbiome*. Evidence shows that many species shift and/or expand their geographical ranges following warming climate; yet, the response of host-associated microbiome communities to species range shifts has received less attention, especially in observational studies. Here, we use two British butterfly species for which detailed long-term distributional data is available, and document for the first time a variety of effects of warming-induced range expansion on the diversity and composition of their microbiome. Our results show similar diversity and species-specific responses in the evenness of the gut microbial communities in the two butterflies. However, butterflies at the expanded ranges showed (i) a larger variability in the microbiome diversity, and (ii) a smaller core community of bacteria that is to a large extent a subset of the bacterial core community in the original range. The microbiome is responsible for many aspects of the host’s physiology and growth, and for ecosystem function, so if the changes in the gut microbial communities reported here apply to other species and taxonomic groups, the potential impact to biodiversity and functioning after range expansion could be severe.

## Introduction

Current climate change takes place at unprecedented rates when compared to historical records (Jones & Mann 2004), and its ecological consequences have been documented for thousands of species worldwide (Parmesan & Yohe 2003). Among the well-documented responses to increasing temperature are the organisms’ range shifts towards the poles and higher altitudes (Grabherr, Gottfried & Pauli 1994; Parmesan 1996; Stange & Ayres 2001; Parmesan & Yohe 2003; Hickling et al 2006; Hoegh-Guldberg et al. 2008; Burrows et al. 2011; Chen et al 2011; Pateman et al. 2012; Pauli et al. 2012; Rassmann et al 2014). There is ample evidence that such range shifts can impact species diversity (Parmesan 1996, 2006), community composition (Wahl et al 2011), body-size structure (Albouy et al 2014), phenotypic plasticity (Matesanz et al. 2010), and species interactions (e.g. Tylianakis et al 2008; Ledger et al 2012, Memmott et al 2007; Durant et al 2007). For example, as species of different trophic levels do not necessarily shift their ranges equally in response to climate change, established food web interactions may become disrupted or spatially decoupled (Phillips et al 2010; Rassmann et al 2014; Gomez-Ruiz & Lacher 2019). These asynchronous responses in space following recent climate change can result in shifting patterns of species dominances and evenness within communities, and potentially to the formation of non-analogue communities (Huntley 1991; Walther 2004; Kullmann 2006; Hobbs et al 2009). Such changes can have important consequences for further population responses to climate change and for ecosystem processes, and therefore need to be considered in making inclusive predictions about the effects of a warming climate on biodiversity and ecosystems (Gilman et al 2010).

Despite the plethora of empirical studies documenting a spatial expansion in populations following climate change, the response of host-associated microbiome communities to species range shifts has received less attention. Most species are a habitat for a relatively diverse community of organisms, including bacteria, fungi and viruses. These communities are of major influence on organismal fitness and population ecology, but until recently it was nearly impossible to study microbial communities at a large scale due to methodological limitations. Metagenomic sequencing has made it possible to identify the microbiome of organic samples, and several studies have investigated how microbial communities of host species changes to increasing temperature. For example, recent research has found climate-induced changes in the microbiome community of several taxonomic groups, including *Sphagnum* mosses (Carrell et al 2019), kelp (Qiu et al 2019), and the gut microbiota of amphibian (Fontaine et al 2018), reptile (Bestion et al 2017) and rodent species (Li et al 2020). Collectively, these studies provide evidence that increasing temperatures can impact the diversity and composition of host microbiomes, with implications to the organism’s physiology and survival, and to ecosystem function. However, while these results come mainly from experimental mesocosms, comparative, observational studies of host-microbiome communities in original and expanded distributions of range-shifting species are scarce (Jones et al 2018; Ramirez et al 2019).

In this context, a fundamental question arises: Are there general patterns in the way the microbiome responds to climate change-driven range shifts of the hosts, or are responses merely idiosyncratic – i.e. each host microbiome responds differently? More specifically, what are the effects of warming-induced range expansion on the diversity and composition of the host’s microbiome? Several hypotheses may explain the effects of climate change on the microbial communities of range-expanding hosts. Species undergoing range expansions may escape from antagonistic interactions such as those from their natural enemies. For example, Engelkes et al. (2008) showed that range-expanding plants were released from shoot and root enemy control, whereas Menéndez et al. (2008) found similar parasitoid richness but lower parasitoid attack rates in the expanded range of a butterfly relative to its original range. If this so-called *enemy release* hypothesis is in place, we would expect microbial communities in the expanded range to be a subset of those in the original range. This hypothesis applies to antagonistic interactions; yet, host-microbiome interactions can also be symbiotic. Conversely, range shifts may also lead to the formation of new interactions in the species’ expanded range margins (e.g. Kullman 2006; Collinge & Ray 2009; Hobbs et al. 2009; Lurgi et al 2012). In other words, a shift in spatial distribution range may result in a parallel shift in species composition of microbial communities. However, it remains largely unknown what is the balance between lost and new species and interactions. Besides, most studies do not consider microbial communities, so we know little about the potential changes in the diversity and composition of the host microbiome following warming-induced range shifts of the host.

British butterflies are excellent focal species to study the ecological consequences of range shifts. Their ecology is exceptionally well known and they are highly sensitive, and respond rapidly to, environmental change. This is because, as other insect groups, butterflies are ectothermic, have high reproductive rates and short generation times, and a high ability to disperse, which lead them to respond more quickly to climate change through range shift than longer-lived and less mobile organisms, e.g. plants. Furthermore, detailed long-term distributional data is available for UK butterflies, which is an invaluable source for research on the effects of climate change on species’ ranges. In a study on European butterflies, 63% had ranges that shifted to the north by 35-240 km during the 20^th^ century, while only 3% have shifted to the south (Parmesan 1999). Of the species with the northern range boundary in the UK, some have shown a dramatic range shift in the past decades. For example, the Brown Argus (*Aricia agestis*) has shifted its UK range margin with more than 100 km in the past 25 years, and the Comma (*Polygonia c-album*) has moved northwards 220 km during the same period (UKBMS, www.ukbms.org). Butterflies are also important model species in food web ecology research, and particularly butterfly-parasitoid interactions are well studied in this context (e.g. Klapwijk & Lewis 2014). To a lesser extent, butterfly-pathogen interactions are used to study ecological disease dynamics (e.g. Bradley & Altizer 2005). Surprisingly though, the butterflies’ gut microbiome has been less studied than that of other insect groups (Engel & Moran 2013; Hammer et al 2014; Yun et al 2014; Coyte et al 2015; Nair & Agashe 2016), which contrasts with the detailed information on butterflies’ distribution patterns.

Here, we investigate how warming-induced range shifts impact gut microbial communities in the expanded range of butterflies. Using two butterfly species as case models, we compare diversity, abundance and composition of butterflies’ gut bacterial communities in original and shifted ranges. Specifically, we address two questions: (1) How does climate change-driven range expansion affect the gut microbial diversity of expanding butterfly species?, and (2) What are the effects of climate-driven range expansion on the abundance patterns and the microbiome composition in the expanded margin of butterflies relative to the original range?

## Material and Methods

### Butterfly species, site selection and field sampling

We used detailed data on the distribution and density of UK butterfly species from the UK Butterfly Monitoring Scheme (UKBMS, www.ukbms.org), in the form of long-term (>30 years) biweekly transect data for numerous sites across the UK. The UKBMS has documented recent range shifts of several UK butterfly species in response to climate change, and allowed us to carefully select the species and sites best suited to address our questions. We sampled two butterfly species that have shown a severe shift of their northern range margin during the past 10-25 years: Brown Argus (*Aricia agestis*), and Speckled Wood (*Pararge aegeria*). We chose sites where abundances of these butterflies have recurrently been high in previous years, thus representing locations where the butterflies are already established, abundant, and can be easily sampled. The same sites were chosen to sample ≥1 butterfly species if they coexisted. For each butterfly, we selected sites that were spread out within the distribution range (both in original and expanded margin) to avoid local effects on the microbiome. This protocol resulted in a total of 14 sites (Figure 1): 8 sites for Brown argus (4 in original, 4 in expanded margin), and 6 sites for Speckled wood (3 in original, 3 in expanded margin). Sampling was conducted in June-September of 2014. Each site was visited several times over the flight period of each butterfly species to account for possible variation in microbial diversity due to environmental conditions and age of the butterflies. The general flight times of the species are: Brown Argus: late July – early September, Speckled Wood: July - early September. For each site, we collected 20 adult individuals per butterfly species to analyse gut microbial diversity, and stored the samples at - 20°C.

**Figure 1.**
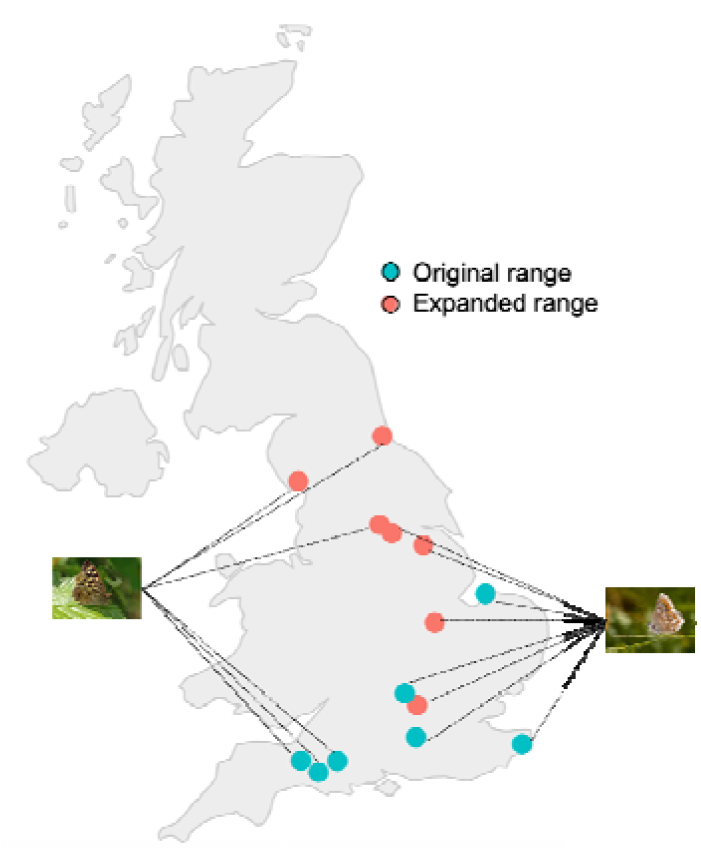
Map of the sampling locations across the distribution range of the two butterfly species. Sites within the expanded range are not always located north of original ranges, despite this is the common case. For each site, 20 adult individuals per butterfly species were collected to analyse gut microbial diversity and composition.

### Metagenomics

After completion of the fieldwork, we extracted genomic DNA from the individual samples in the laboratory. We focus on the gut microbiome because the likelihood of ‘contamination’ is lower than that of external bacteria. We used Illumina High-seq to sequence the bacterial 16S rRNA and fungal ITS ribosomal region of the pooled population samples to determine the bacterial and fungal metagenome for each butterfly species and site. The primers were designed with a population-specific barcode sequence so that amplification products from the different populations could be pooled together for a single library preparation and sequenced on a single lane. Metagenomics sequencing was done at the University of Bristol’s Genomics Facility (http://www.bristol.ac.uk/biology/genomics-facility/). The resulting short reads were assembled into longer contigs, and the microbial taxonomy was determined by comparing the sequences against online databases such as the European Ribosomal RNA database. This protocol allowed us to identify the gut microbiome for each butterfly species – operational taxonomic unit (OTU) - and each site.

### Statistical and network analysis

We calculated various metrics describing the diversity, abundance and composition of microbial communities of butterflies in original *versus* expanded ranges. All analyses were performed in R software (R version 4.0.5; R Core Team, 2020).

#### Diversity and abundance

We measured the number of OTUs (mean ± sd) and different features of OTU abundance distributions. Changes in biological communities by anthropogenic stressors, including climate change, not only affects the number of species, but also their relative abundance and thereby dominance or evenness patterns in most ecosystems (Hillebrand et al 2011). More importantly, evenness often responds more rapidly to environmental constraints than species richness (Chapin et al 2000). Thus, the distribution and variance of species within a host may be more important for aggregate performance of microbial communities than the number of species. To assess how evenly OTU abundance is distributed between original and expanded sites, we fitted *Zipf* models to rank-abundance distributions (RADs). The *Zipf* model is given by the power law *a_r_* = *Np*_1_*r^γ^*, where *a_r_* is the predicted abundance of the OTU rank *r*; *N* is the total number of individuals; *p*_1_ is the estimated proportion of the most abundant OTU (rank 1); and γ is the estimated exponent of the power law (γ<0). Thus, a smaller value of |γ| indicates a more even distribution of abundances. We also estimated *β*-diversity of OTUs (Whittaker’s index) to investigate changes in OTU composition between the butterflies’ original and expanded ranges.

#### Core communities

Ecological communities, including host-associated microbial communities, comprise different types of microorganisms according to the intimacy and repeatability of the association with their host, forming a continuum from core through transient to opportunistic organisms (Schmitt et al 2011; Phalnikar et al 2019). This has prompted the development of the ‘core microbiome’ concept as a way to identify organisms that are consistently found within a particular host species, likely performing core metabolic functions (Astudillo-Garcia et al 2017; McDevitt-Irwin et al 2017; Lurgi et al 2019). To examine the structuring of the most prevalent interactions between butterflies and bacteria that are in close association with them, and whether they change following range expansion between original and shifted ranges, we constructed the butterfly core–microbiome community. The butterfly core–microbiome community was defined as the subset of OTUs found in butterfly hosts that are (i) present in at least two-thirds of the sites in each part of the butterfly’s range, and (ii) present with a relative abundance ≥0.001% across the whole dataset within each part of the range (Astudillo-García et al 2017; Lurgi et al 2019). Results were relatively robust to changing abundance criteria (see Table S1 in Supplementary Material).

## Results

### Diversity and abundance of butterfly’s microbiome

Richness of microbial communities varies across different host species. As with most organisms inhabiting the guts of most insect species, bacterial species comprise all or the majority of organisms of the gut microbiome. With respect to bacterial taxonomy, our findings are generally congruent with previous work on insect-associated bacterial communities (Engel & Moran 2013; Yun et al 2014; van Schooten et al 2018). The most abundant and frequent OTUs in the studied butterflies are also common members of other insect-associated microbiomes: *Proteobacteria* and *Firmicutes* were the dominant phyla and represented a 50% and 21% of the total sequences, respectively, with abundant phyla of *Bacteroidetes, Actinobacteria*, and *Tenericutes* in some butterfly samples (Figure S1 and Table S3 in Supplementary Material). There was no evidence to support differences in the mean number of OTUs for butterflies at the expanded range (*F* = 1.472, *p* = 0.248), suggesting that range expansion did not result in a net gain or loss of OTUs. Yet, a larger variability in the number of OTUs was observed in expanded versus original ranges (Brown argus: {31, 55} vs {39, 53}; Speckled wood: {31, 46} vs {39, 45}) (Figure 2).

**Figure 2.**
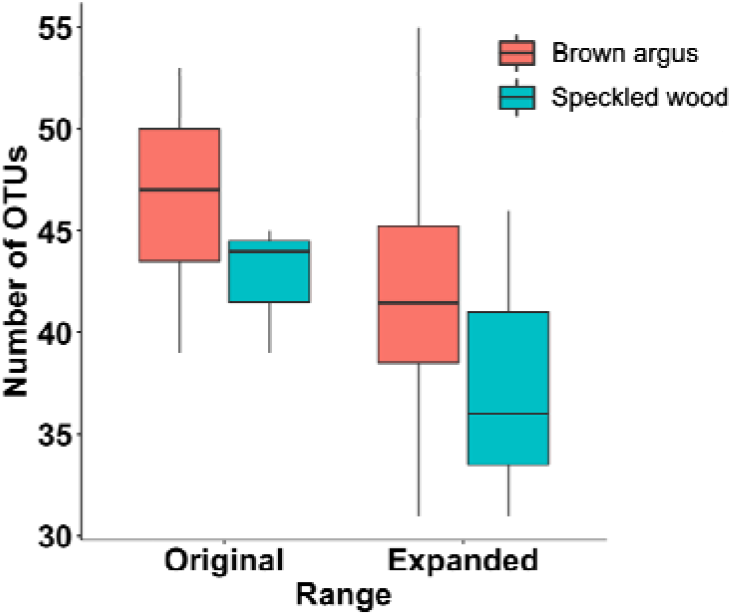
Diversity of the butterflies’ gut microbiome in original and expanded ranges. Plots show the mean and standard deviation of the number of bacterial OTUs for per butterfly species. There was no evidence supporting differences in mean OTU richness (F = 1.472, p = 0.248), despite a larger variability in expanded versus original range is clear.

RAD analysis shows similar and more even distribution of OTU abundances in the butterflies’ original range for brown argus and speckled wood, respectively (*Zipf* fits, mean |γ| across sites: Brown argus = {0.605 (original), 0.6 (expanded)}, Speckled wood = {0.61 (original), 0.67 (expanded)}). That is, whereas no difference in evenness are observed for brown argus, certain bacteria are more dominant in the expanded margins of the speckled wood populations (Figure 3). When divided into abundance categories we find that, whereas the abundance of dominant bacteria is similar in both original and expanded ranges, differences in the bacterial abundance distribution mostly arise at intermediate and low levels of abundance (Figure 4). This pattern is consistent either aggregating all sites per butterfly or considering individual sites separately (Figure S2 in Supplementary Material). This is further supported by the *β*-diversity analysis of the three abundance categories (>0.01, 0.01 – 0.001, <0.001): despite no major turnover is observed within the most abundant bacteria, intermediate and rare bacteria show a high turnover (Whittaker’s index for most, intermediate and least abundant OTUs: Brown argus = {0.22, 0.34, 0.75}, Speckled wood = {0.39, 0.48, 0.71}). Although these numbers can slightly vary when using different thresholds to establish abundance categories (Table S2 in Supplementary Material), *β*-diversity is always lower for the dominant bacteria and increases as bacteria become less abundant.

**Figure 3.**
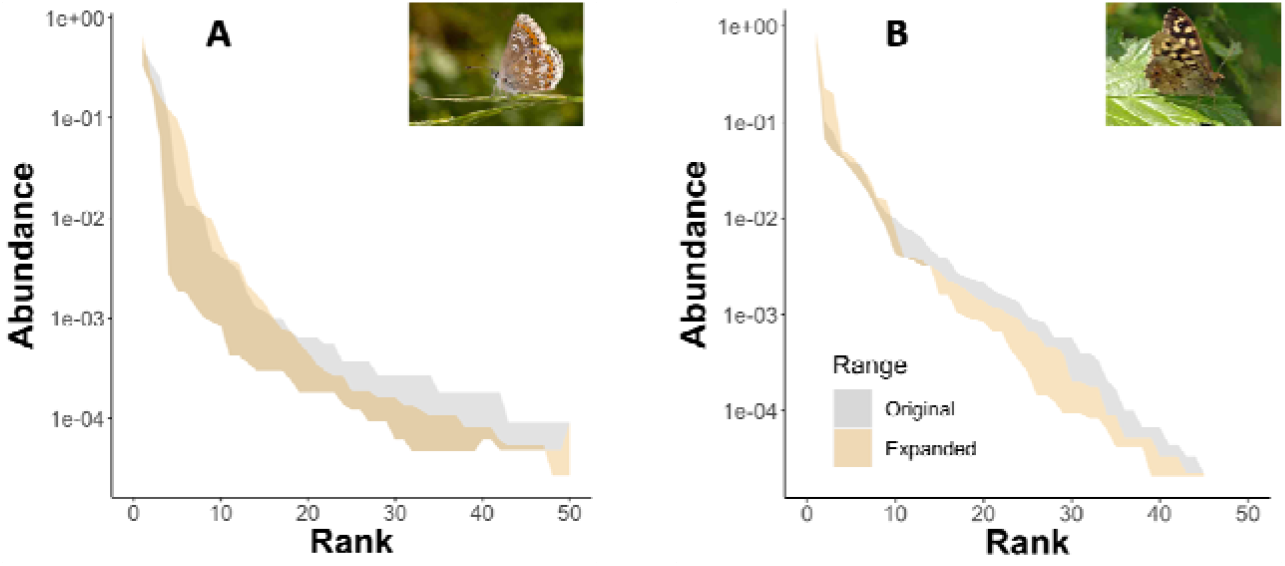
Rank-abundance distribution of bacterial OTUs in original and expanded ranges. **(A)** Brown argus; **(B)** Speckled wood. Abundance values corresponding to all sites within each range are contained within the shadows, whose limits are determined by the minimum and maximum values across the range of bacteria abundances within each range (original and expanded)

**Figure 4.**
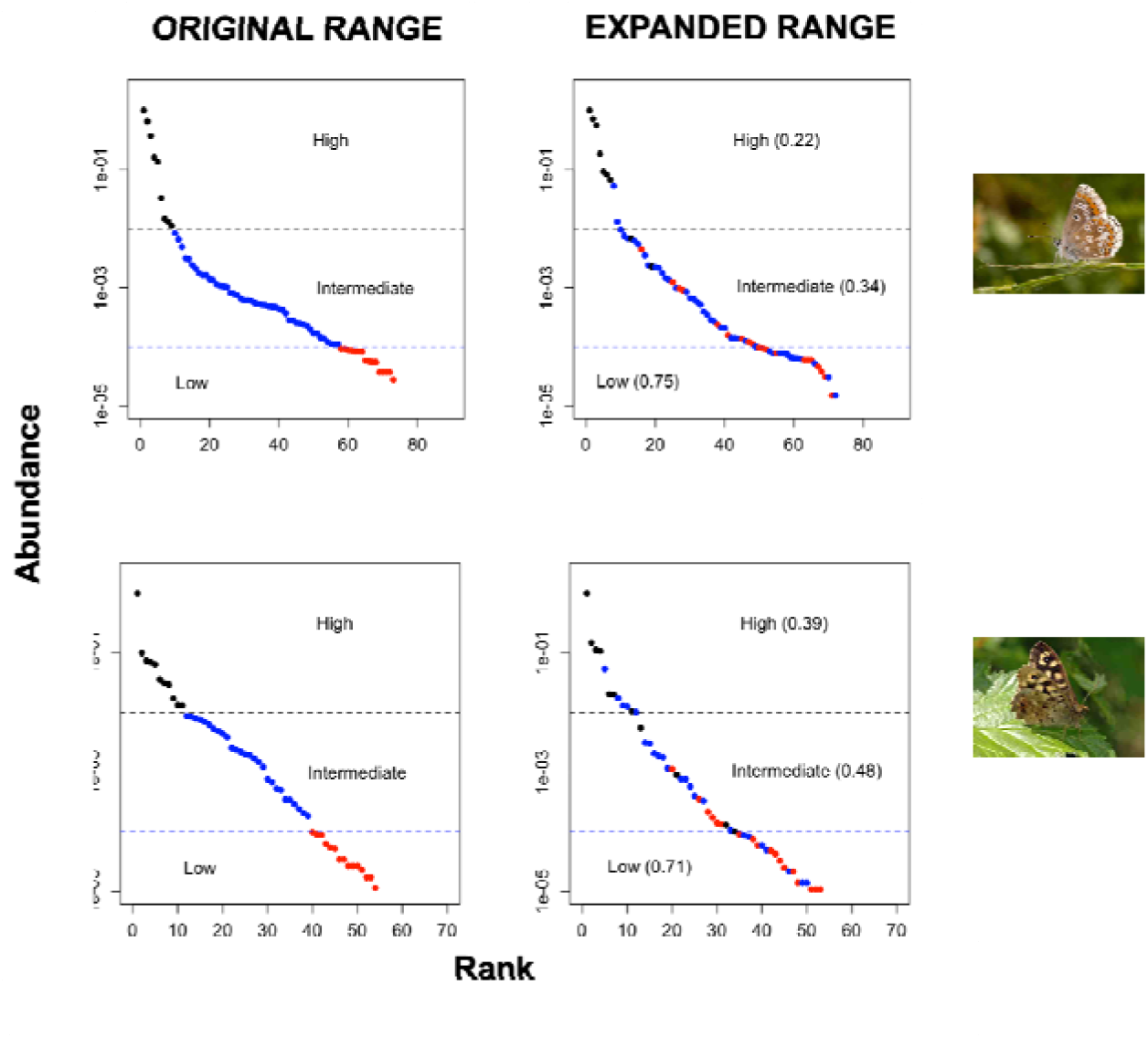
Rank-abundance distribution of bacterial OTUs, divided into abundance categories (black – high, blue – intermediate, red – low), in original and expanded ranges. Upper row: Brown argus; Bottom row: Speckled wood. Abundance categories (and their colors) are established for bacteria within the original range and maintained in the expanded range graphs, allowing to visualize bacterial turnover between original and expanded ranges. Most of the differences in the abundance distribution of bacteria arise at intermediate and low levels of abundance, whereas dominant bacteria are persistent across original and expanded ranges. Numbers indicate β-diversity (Whittaker’s index) for the different abundance categories (higher βindicates higher turnover).

### Core communities

Figure 5 shows the core communities of each butterfly species with original and expanded range sites. Despite the application of such strict constraints on the definition of the core microbiome (see Material and Methods), a relatively large butterfly core–microbiome community remained. Butterflies in the original sites have a larger core community of gut bacteria relative to that of the expanded regions. Besides, the core community at the expanded margin is, to a large extent, contained within the original range for the two butterflies, thus suggesting that butterflies left some bacteria behind during climate-driven range expansion. The butterfly-microbiome also acquires new bacteria at the expanded margin, this resulting in a similar mean number of OTUs (Figure 2). These results were robust to changing abundance criteria (see Table S1 in Supplementary Material): for both butterflies, at least two-thirds of the core community of bacteria at the expanded range are a subset of the core community within the original range of distribution.

**Figure 5.**
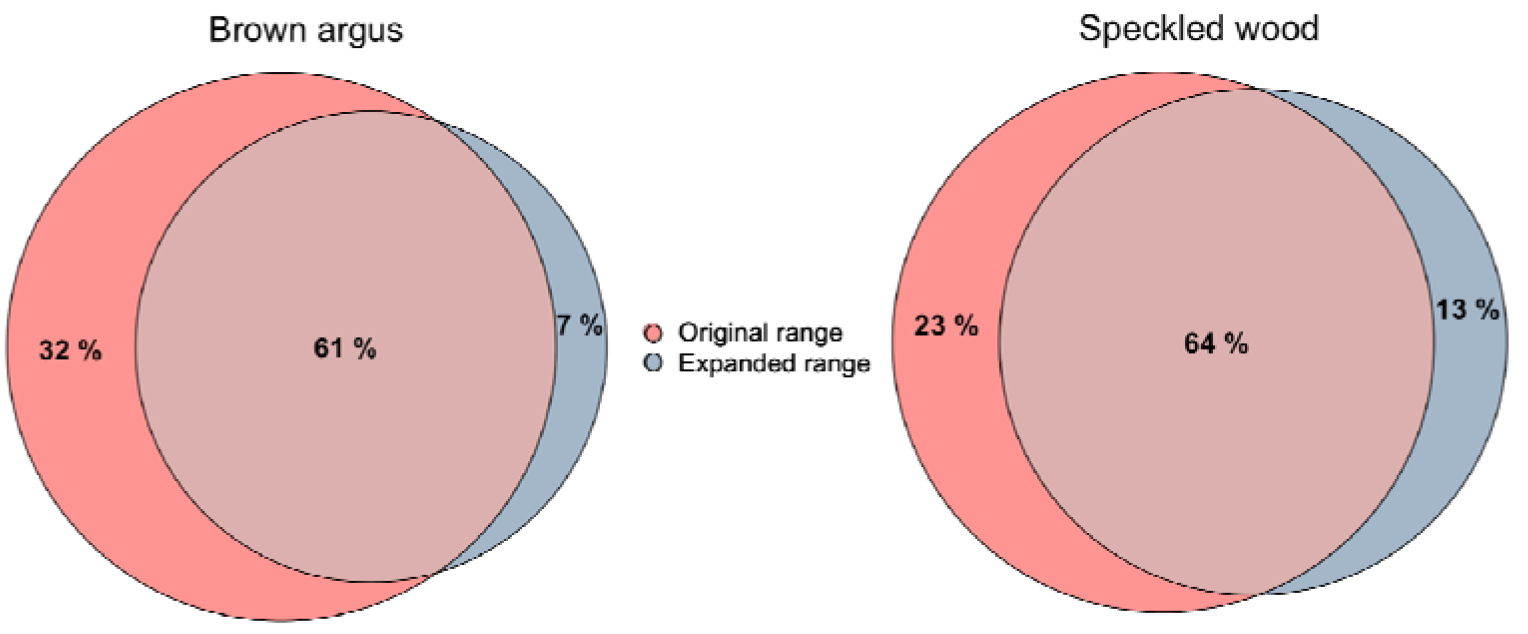
Butterfly-bacteria core communities across distributional ranges. For each butterfly, original and expanded range sites are included. The number of bacterial OTUs in original and expanded range sites are indicated, as well as the number of shared bacterial OTUs across the entire distribution range of each butterfly species.

## Discussion

To our knowledge, this is the first observational study in real-world communities that investigates changes in the diversity and composition of the host-microbiome in species that have experienced warming-induced range expansion. Using two butterfly species as case studies, our results reveal changes in butterfly microbial communities in the expanded range of the butterflies’ distribution. Despite no significant changes in mean OTU numbers and species-specific responses in evenness, two general patterns were observed at the shifted ranges of the butterflies that are consistent across the two species: (i) a larger variability in the diversity of gut bacterial communities, and (ii) a smaller core community of bacteria that is to a large extent a subset of the bacterial core community in the original range.

Despite our results find support for negative effects of range shifts on the average bacterial diversity of the gut microbiome (e.g. core communities are smaller at expanded ranges), they are compensated with the acquisition of new bacteria. The balance between lost and gained bacteria is neutral, thus resulting in microbiomes that are not significantly different in terms of mean OTUs despite a larger variability in their numbers. Our analysis also reveals that the core bacterial microbiome of the butterflies becomes less diverse after range expansion. Moreover, the core microbiome at the expanded margin is, to a large extent, contained within the original range for both butterflies. Such reduction in the diversity of the core bacterial communities in parallel to similar mean, but higher diversity levels, may be driven by several, non-exclusive factors. First, butterflies acquire their microbiome from the environment via food sources (horizontal transmission), and thus smaller core communities at the expanded ranges of the butterflies’ distribution would result from the fact that only a fraction of the bacteria found in the butterflies’ core gut microbiome is common in both parts of the distribution range. However, range-shifting populations tend to have broader diets, and this may counteract the effects of potentially lower diversity of food sources that provide core bacteria (Braschler & Hill 2007; Pateman et al 2012; Lancaster 2016; Eloy de Amorim et al 2017; Suehiro et al 2017; Shively et al 2018; Singer & Parmesan 2020); indeed, Lancaster (2020) analyzed 4410 butterfly species across the globe and found that geographic range expansions are linked to butterflies’ diet expansion. Collectively, our results suggest that smaller core microbiomes in the expanded range of the butterflies’ distribution are compensated by broader diet breadths, this leading to bacterial communities that are highly variable, yet similar in their mean OTU diversity.

An alternative explanation to the larger variability in bacterial diversity is that butterflies at the expanded range are going through a transient state. On one hand, the gut microbiome is horizontally transmitted and depends on both the availability of food resources and the butterflies’ diet breadth. Diet breadth tends to increase with latitude and decline with population age in butterflies (Singer & Parmesan 2020). Thus, the high variability of bacterial diversity observed in the expanded ranges could reflect a transient state where the studied butterflies’ populations are still young and not yet stabilised. Such variability could also be indicative of the strength of host-bacterial interactions; in the case of symbiont interactions, low variability would indicate more obligate interactions between butterflies and bacteria (Phalnikar et al 2019). On the other hand, it could also be argued that the studied butterflies have already reached an equilibrium state in their expanded margins, as (i) butterflies exhibit a strong response to warming given their ectothermic nature, short generation times, and high dispersal ability, and (ii) we chose sites where the butterflies were already present in the last 10-25 years and with high abundances in previous years before sampling. However, the studied butterflies have 1-2 generations per year, so it is debatable whether their populations are at equilibrium. Further, newer populations in the expanded margins are usually smaller and/or more isolated, and this may reduce microbial exchange via the environment. Studies that include temporal resolution will be needed to determine whether the observed reduction in microbiome diversity reflects a transient state.

Our results show that bacteria richness and evenness can respond independently to warming-induced range expansion, as shown by changes in the bacterial abundance distributions of one of the butterflies’ microbiome. Specifically, whereas no difference is reported for brown argus, bacterial abundances in the shifted ranges of speckled wood populations are less evenly distributed compared to the original range. This generally agrees with recent findings documenting that species richness and abundances can respond simultaneously to global change factors in various and independent ways (e.g. Chapin et al 2000; Hillebrand et al 2011; McWilliams et al 2019). Our results further show that the identity of the most dominant bacteria in the microbiome persists in both parts of the butterflies’ range, and that the effects of range expansion on the butterflies’ microbiome has a higher impact in bacteria with intermediate and low levels of abundance. These are the less stable bacteria that experience the highest turnover between original and shifted ranges. Differences in the abundance distributions and evenness patterns of bacteria could be explained by the fact that, in butterflies, the impact of gut microbes on their insect hosts range along a continuum from strong to weak dependence, to no association (Phalnikar et al 2019). Moreover, even when acquired independently each generation, insect gut communities are not expected to be random assemblages of bacteria from the food or local environment (most studies show differences between microbes in the guts *vs* those ingested with food). Rather, gut communities seem to be dominated by widely distributed bacteria that appear to colonize hosts opportunistically (Engel & Moran 2013). Thus, whereas rare bacteria might be more responsive to warming and/or less widely distributed, abundant gut bacteria become abundant because they establish intimate associations with their host butterflies (e.g. symbiosis) and/or are present in high abundances across the distribution range of butterflies (opportunistic).

How do the observed changes in the microbiome diversity and composition affect butterflies’ survival and ecosystem function? Although this is beyond the scope of this work, the results here reported have several conservation implications. First, extinction risk decreases with evenness, as shown by e.g. experiments where plant extinctions were higher in plots sown with low compared to high evenness (Wilsey & Polley 2004). In the case of speckled wood populations, this suggests that butterflies’ microbiome at the shifted ranges may lose more microbial diversity, especially that of rare bacteria. Second, experimental studies across different taxonomic groups show that alterations in diversity and composition of host microbiomes in response to warming can strongly impact the host’s physiology and survival (Bestion et al 2017; Fontaine et al 2018; Carrel et al 2019; Qiu et al 2019; Li et al 2020). Besides, changes in evenness are as important as changes in species richness and can have important consequences for ecosystem function (Hillebrand et al 2011). Third, our analyses did not discriminate between antagonistic and symbiotic bacteria, so relative changes in the proportion of these interactions might be taking place. In contrast to the benefits associated with the release from antagonistic interactions (e.g. predators, pathogens) on range-expanding species (Callaway et al 2004; Mueller & Hellmann 2008; Mitchell et al 2006; Lavergne et al 2010; Hellmann et al 2012), a higher symbiont gut bacterial diversity is often beneficial to hosts (Bolnick et al 2014). Thus, climate-driven changes in bacterial composition could be detrimental for host butterflies, particularly if it entails a loss of essential functions (Mandrioli 2012; Le Chatelier et al 2013; Kikuchi et al 2016).

Our study has several limitations that future analyses should overcome. For example, we used two butterfly species as case studies. Although these butterflies were selected because of the availability of detailed information on their range expansion, adding more species and sites would provide a more complete picture of the effects of warming-induced range expansion and whether the results here reported can be generalized to other butterflies and different taxonomic groups. Further, despite each site was visited several times over the butterflies’ flight period to account for possible variation in microbial diversity, we acknowledge that this procedure captures within-year variation but does not account for the long-term patterns of range expansion. Studies should incorporate a dynamical perspective with time-series to overcome this limitation and reveal whether range-shifting butterflies are going through a transient state or have already reached a dynamical equilibrium within their expanded ranges. Despite these limitations, to our knowledge this is the first comparative, observational study of host-microbiome communities in original and expanded distributions of range-shifting species, and a useful first step to contribute to furthering our understanding of the effects of warming on the diversity and composition and of host bacterial communities after range expansion.

There is ample evidence across geographical areas and taxonomic groups on species’ range shifts towards the poles and higher altitudes as a response to warming climate. Here, we have used two butterfly species as case studies and documented for the first time a variety of effects of warming-induced range expansion on the diversity and composition of their gut microbial communities. These range from non-significant changes (bacterial richness), to idiosyncratic or butterfly-dependent changes (evenness patterns), and changes that are consistent across the two butterflies (variability in bacterial richness, core microbiome composition). The microbiome is responsible for many aspects of the host’s physiology and growth, and for ecosystem function, so if the changes in the gut microbial communities reported here apply to other species and taxonomic groups, the potential impact to biodiversity and functioning after range expansion could be severe. Future studies should aim at investigating the degree to which the patterns observed in this study are consistent and the consequences of its alteration to species and ecosystems.

## Acknowledgements

We thank Emmeline Topp, Joanne Morten, Kees Wanders, Alicia Hodson and Jon Bridle for their assistance in conducting field and lab work, and Tom Batstone for the bioinformatic analyses. This research was financially supported by a British Ecological Society Large Research Grant.

## This file includes

Tables S1 and S3

Figures S1 and S2

**Table S1.** Butterfly’s core-microbiome community composition in expanded margins. The butterfly core–microbiome community is defined as the subset of OTUs found in butterfly hosts that are (i) present in at least two-thirds of the sites in each part of the butterfly’s range, and (ii) present with a relative abundance ≥0.001% across the whole dataset within each part of the range. The application of stricter criteria did not significantly change the original results. Numbers represent the percentage of the microbiome in the expanded margins that is contained in the original range of the butterflies.

|  | **Presence $\geq 2/3$** |  |  | **Abundance $\geq 0.001\%$** |  |
| --- | --- | --- | --- | --- | --- |
|  | *Abundance $\geq 0.001$* | *Abundance $\geq 0.01$* | *Abundance $\geq 0.1$* | *Presence = 100%* | *Presence $\geq 2/3$* |
| **Brown argus** | 0,67 | 0,75 | 1 | 0,69 | 0,80 |
| **Speckled wood** | 0,83 | 0,625 | 1 | 0,88 | 0,79 |

**Table S2.**
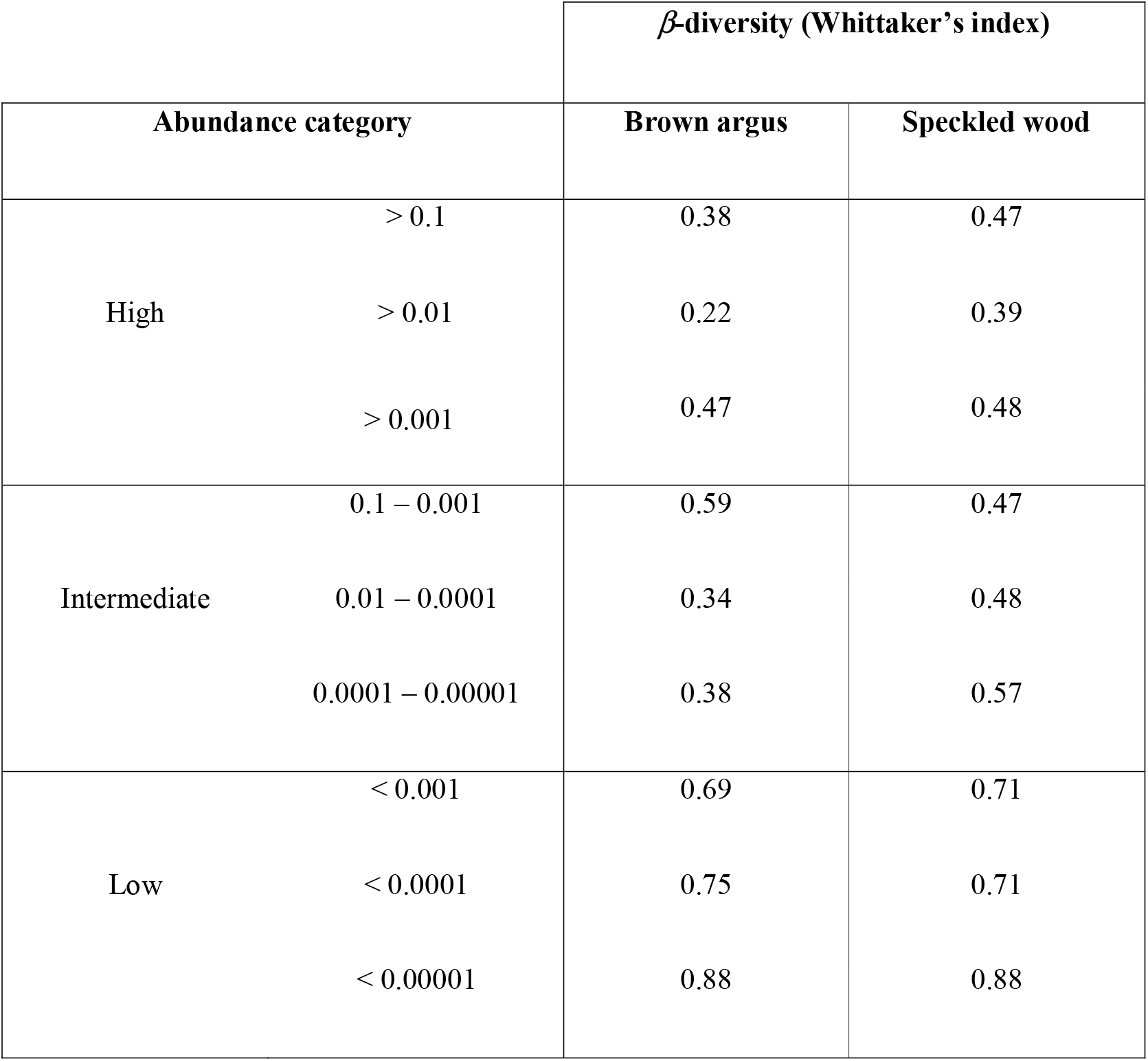
*β*-diversity, measured as Whittaker’s index, for different abundance categories. For any abundance category, *β*-diversity is always lower for the dominant bacteria and increases as bacteria become less abundant.

**Table S3.**
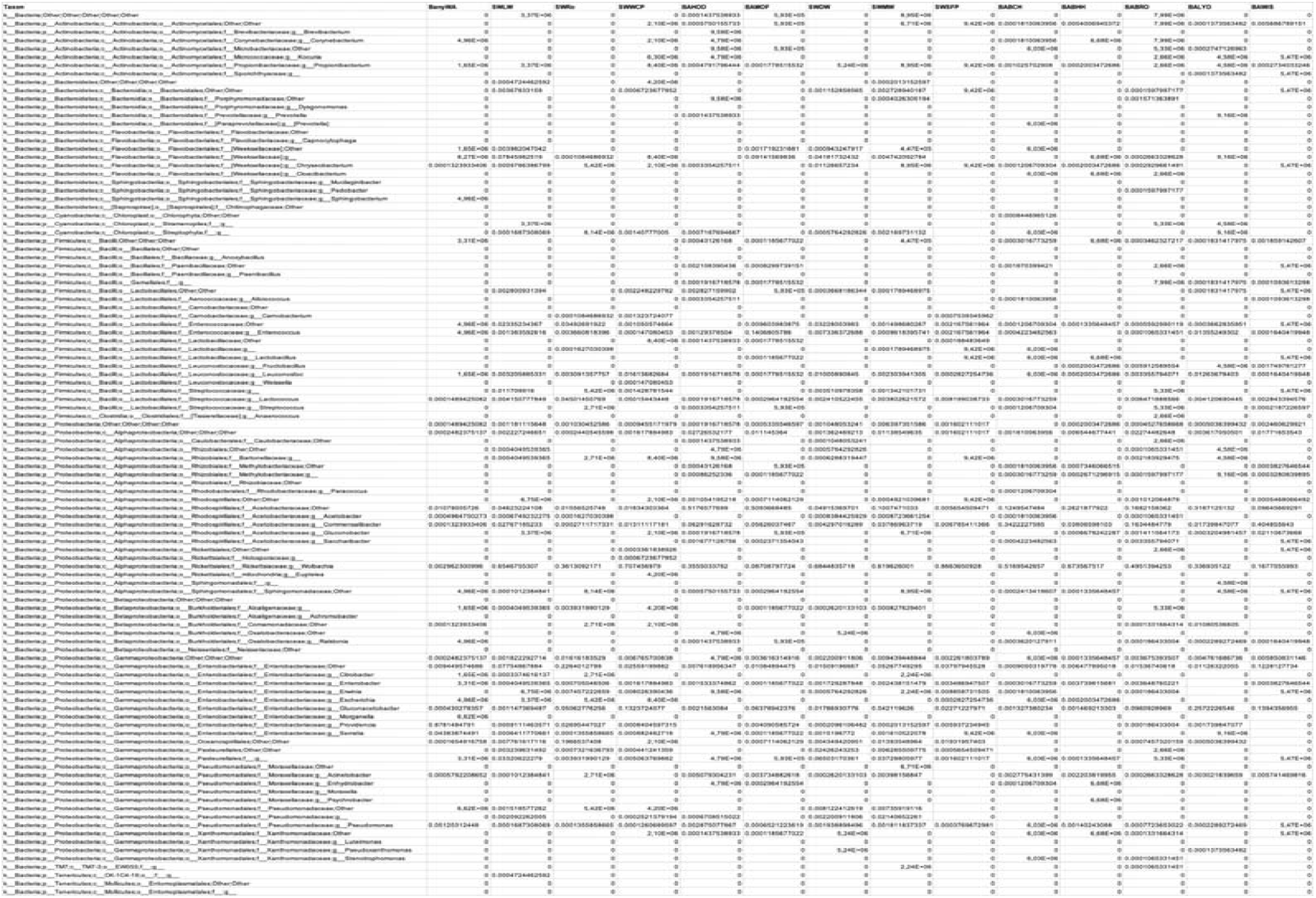
Taxa (OUT) identification for each sampling site. BA = Brown argus; SW = Speckled wood.

**Figure S1.**
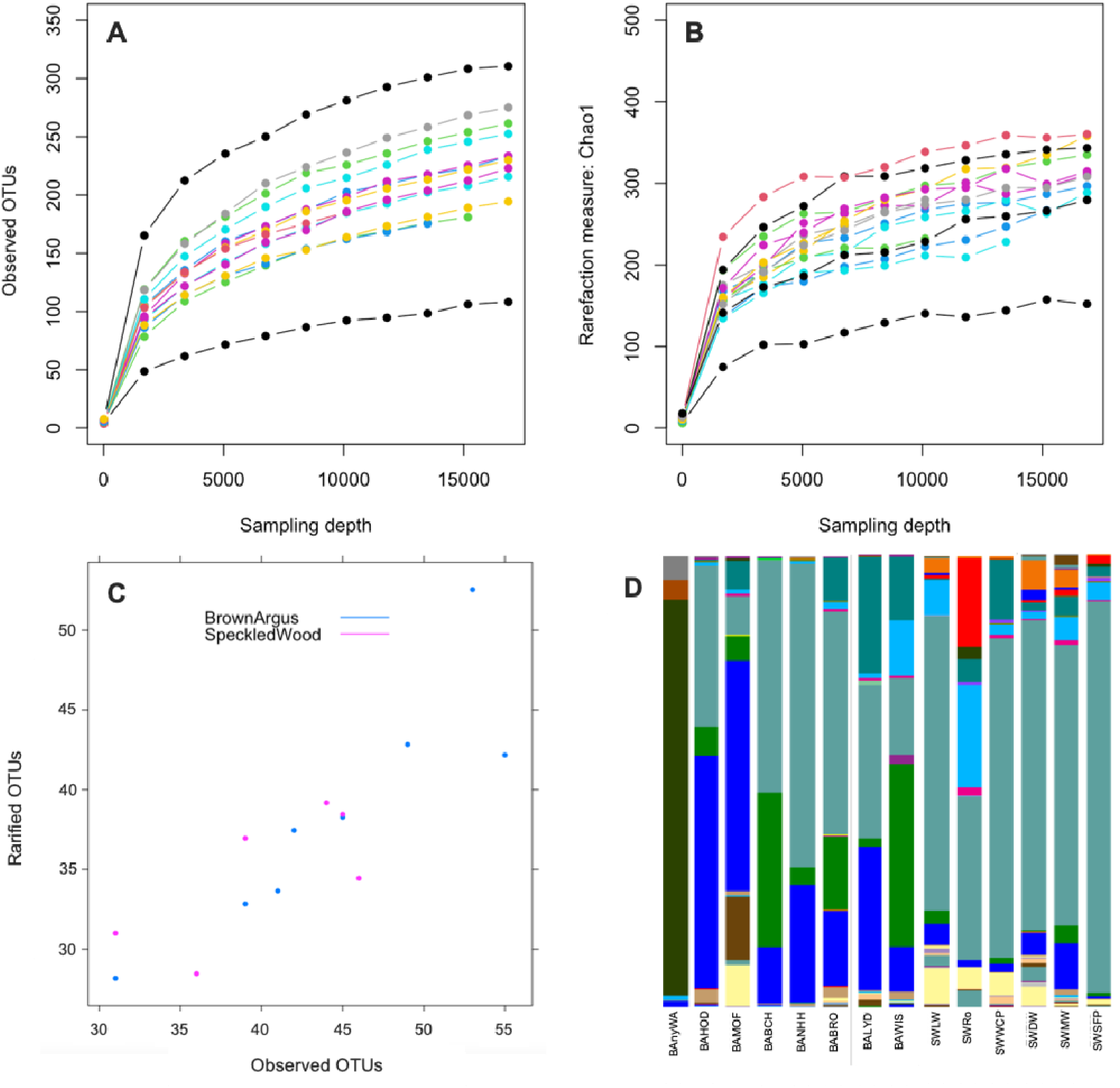
Rarefaction curves described for **(A)** Observed OTUs and **(B)** Chao1 metric among all sample sites. **(C)** Correlation between observed OTUs and rarefied OTUs. **(D)** Relative abundance of OTUs for each sampling site (see table S3 for taxa ID).

**Figure S2.**
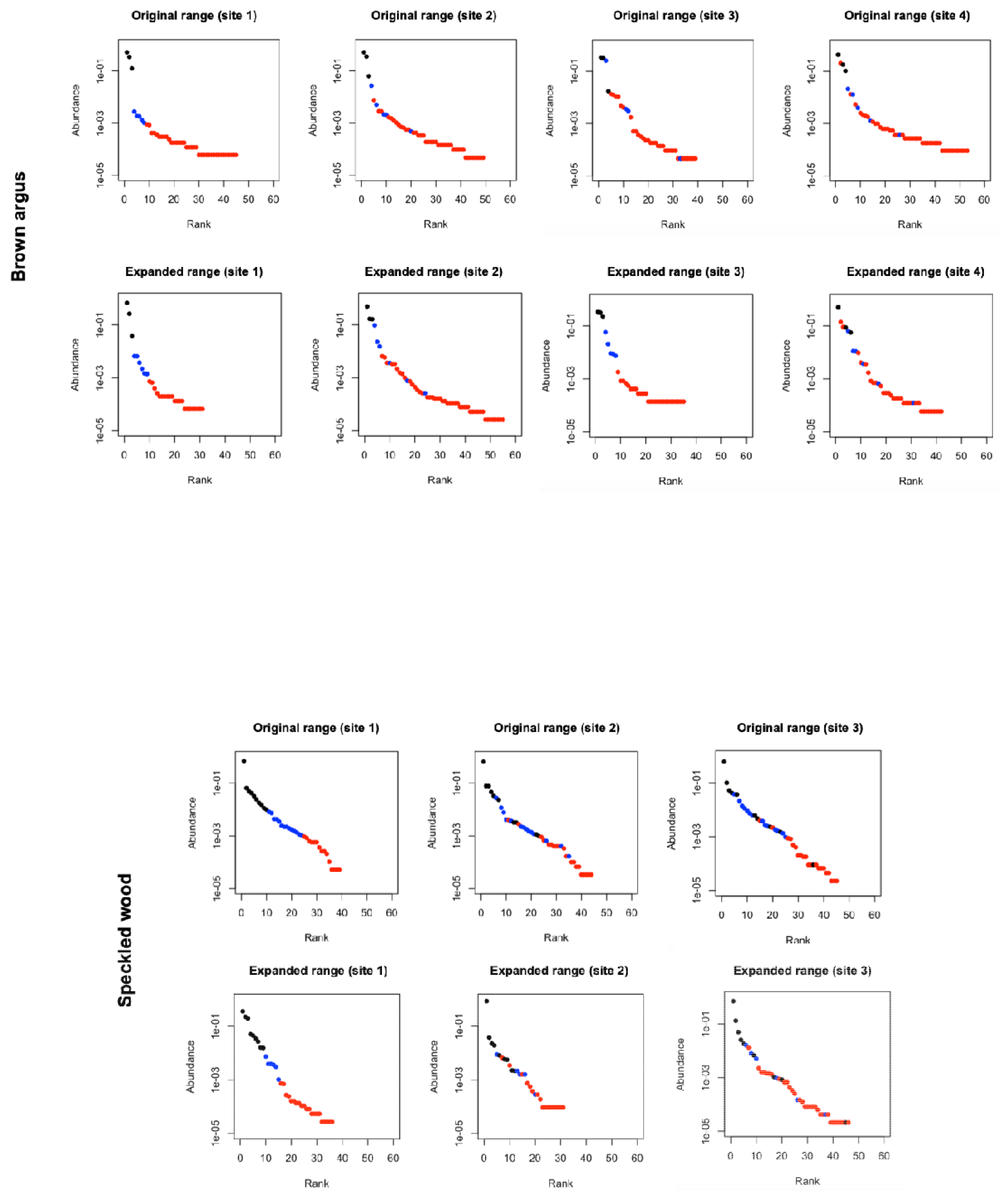
Rank abundance distributions of butterflies’ microbial communities per sampling site. The observed patterns are generally consistent to those documented when pooling all sites together (main text).

